# Supraphysiological protection from replication stress does not extend mammalian lifespan

**DOI:** 10.1101/2020.01.17.910133

**Authors:** Eliene Albers, Alexandra Avram, Mauro Sbroggio, Oscar Fernandez-Capetillo, Andres J Lopez-Contreras

**Affiliations:** Department of Cellular and Molecular Medicine, Center for Chromosome Stability and Center for Healthy Aging, University of Copenhagen, Copenhagen, Denmark; Genomic Instability Group, Spanish National Cancer Research Centre (CNIO), Madrid, Spain

## Abstract

Replication Stress (RS) is a type of DNA damage generated at the replication fork, characterized by single-stranded DNA (ssDNA) accumulation, and which can be caused by a variety of factors. Previous studies have reported elevated RS levels in aged cells. In addition, mouse models with a deficient RS response show accelerated aging. However, the relevance of endogenous or physiological RS, compared to other sources of genomic instability, for the normal onset of aging is unknown. We have performed long term survival studies of transgenic mice with extra copies of the *Chk1* and/or *Rrm2* genes, which we previously showed extend the lifespan of a progeroid ATR-hypomorphic model suffering from high levels of RS. In contrast to their effect in the context of progeria, the lifespan of *Chk1, Rrm2* and *Chk1/Rrm2* transgenic mice was similar to WT littermates in physiological settings. Most mice studied died due to tumors -mainly lymphomas-irrespective of their genetic background. Interestingly, a slightly higher percentage of transgenic mice developed tumors compared to WT mice. Our results indicate that supraphysiological protection from RS does not extend lifespan, indicating that RS may not be a relevant source of genomic instability on the onset of “normal” aging.

## Introduction

For decades, researchers have investigated the process of aging: a decline in health over time, which is generally considered to be due to an accumulation of cellular damage [1,2]. This cellular damage has different causes, but genomic instability is considered one of the main factors that contribute to cellular aging [3]. Genetic damage such as point mutations, chromosomal rearrangements, aneuploidy and copy number variations accumulate during life and are caused by endogenous and exogenous sources, including UV radiation, ionizing radiation and reactive oxygen species (ROS) [3,4]. While the accumulation of genomic instability over time can cause cellular senescence and apoptosis in the case of aging, in other cases it can lead to uncontrolled cellular proliferation, and is therefore associated with an increased cancer risk [3,4].

Defects in DNA repair proteins lead to an accumulation of DNA damage and thus contribute to accelerated aging, as is the case for several human diseases including Cockayne, Bloom and Werner syndromes, trichothiodystrophy and ataxia telangiectasia [5,6]. In addition, several mouse models have confirmed that mutations in DNA repair proteins lead to accelerated aging. These mice exhibit accelerated aging and aging-related phenotypes such as alopecia, grey hair, osteoporosis, cachexia, neurological abnormalities, retinal degeneration and a predisposition to a wide variety of cancers [5,7–12]. Taken together, this indicates that an exacerbated accumulation of DNA damage leads to premature aging, albeit the actual contribution of DNA damage to normal aging remains to be elucidated. More importantly, which types of DNA damage play a central role in the context of aging is still unknown.

Besides strategies to accelerate aging, genetic manipulation studies in worms, flies and mice have also successfully lead to increases in lifespan, suggesting a direct implication of the manipulated genes in normal aging [1,13]. For instance, mice overexpressing the spindle assembly checkpoint protein BubR1 show a reduction in age-dependent aneuploidy, reduced incidence of cancer, and an increased healthy lifespan compared to WT mice [14]. Another relevant mouse model with delayed aging is a mouse model with extra copies of tumor suppressor genes *Trp53* and its positive regulator *Arf*, which in addition to having an increased median lifespan has a reduced tumor incidence [15]. The median survival is increased further in mice with constitutive telomerase reverse transcriptase (TERT) overexpression in addition to extra copies of *Trp53* and *Arf* [16].

In recent years, replication stress (RS) has been acknowledged as an important source of endogenous DNA damage [17]. RS is a type of DNA damage that occurs when obstacles to replication lead to an accumulation of single stranded DNA (ssDNA) at stalled replication forks, which is recognized by ssDNA binding protein RPA. This initiates a signaling cascade involving Ataxia Telangiectasia and Rad3-related (ATR) kinase and CHK1 which promotes DNA repair, cell cycle arrest, and apoptosis [18–20]. Similar to other types of DNA damage, RS has been linked to aging. For instance, aged hematopoietic stem cells (HSCs) exhibit increased levels of RS compared to young HSCs [21]. In addition, mutations in the ATR gene cause Seckel syndrome in humans, which is characterized by progeria, growth retardation, microcephaly, mental retardation and dwarfism [22](OMIM210600). The involvement of RS in premature aging has also been shown experimentally with a mouse model for Seckel syndrome [12]. ATR-Seckel mice exhibit a phenotype similar to that of human patients, which is further aggravated in combination with several cancer-driving mutations such as the *Myc* oncogene or the absence of the tumor suppressor p53 [12,23]. ATR-Seckel mice show high levels of RS during embryonic development, accelerated aging in adult life and early lethality [12]. Interestingly, mice harbouring extra alleles of CHK1 (*Chk1*^Tg^) or of the ribonucleotide reductase (RNR) regulatory subunit RRM2 (*Rrm2*^Tg^), which is a limiting factor for dNTP production, improved the lifespan and alleviated the progeroid phenotype of ATR mutant mice [24,25]. These *Chk1* and *Rrm2* transgenic mice carry bacterial artificial chromosome (BAC) alleles of the respective genes, including exons and intron, under their own endogenous promoters. This strategy provides supraphysiological levels of these factors while preventing overexpression in tissues where these genes are normally not expressed, and was proven successful with the *Trp53* BAC-transgenic mouse model [26]. Collectivelly, these studies suggested that RS might have important implications in mammalian aging. However, the effect of *Chk1* and *Rrm2* expression levels on normal aging, in mice with physiological levels of ATR, remains to be elucidated.

In the current study, we investigated the effect of supraphysiological levels of CHK1 and RRM2, which confer extra protection against RS, on normal aging. We utilized cohorts of WT, *Chk1*^Tg^, *Rrm2*^Tg^ and *Chk1*^Tg^;*Rrm2*^Tg^ mice to asses tumor-free survival of these mice. We found no differences in survival between the genotypes and all mice exhibited similar signs of aging, although there was a slightly higher tumor incidence in the transgenic mice compared to WT mice. In the light of these data, we propose that endogenous RS is not a major source of DNA damage contributing to normal aging.

## Results

### Generation of mice with supraphysiological levels of CHK1 and RRM2

In order to investigate the effect of RS on lifespan, we used the *Chk1*^Tg^ and *Rrm2*^Tg^ mouse models previously generated in our laboratory [24,25]. *Chk1*^Tg^ and *Rrm2*^Tg^ mice were crossed in order to obtain *Chk1*^Tg^, *Rrm2*^Tg^ and *Chk1*^Tg^;*Rrm2*^Tg^ mice. Transgenic mice were not phenotypically distinguishable from WT littermates and were born in accordance with Mendelian ratios (Chi-square p-value = 0.243) (Table 1).

**Table 1:** offspring from C*hk1*^Tg^ x *Rrm2*^Tg^ and *Chk1*^Tg^;*Rrm2*^Tg^ x WT crosses.

|  | WT | *Chk1*<sup>Tg</sup> | *Rrm2*<sup>Tg</sup> | *Chk1*<sup>Tg</sup>;*Rrm2*<sup>Tg</sup> |
| --- | --- | --- | --- | --- |
| Observed | 84 | 66 | 91 | 83 |
| Expected | 81 | 81 | 81 | 81 |

### MEFs with increased levels of RRM2 and CHK1 are protected against RS

To confirm that supraphysiological levels of CHK1 and RRM2 could protect cells against RS, we generated mouse embryonic fibroblasts (MEFs) from crosses between *Chk1*^Tg^ and *Rrm2*^Tg^ mice. *Chk1*^Tg^ and *Rrm2*^Tg^ MEFs have been previously described to be resistant to RS induced by the ribonucleotide reductase inhibitor hydroxyurea (HU) [24,25]. Genomic PCR genotyping confirmed the four different MEF genotypes: WT, *Chk1*^Tg^, *Rrm2*^Tg^ and *Chk1*^Tg^;*Rrm2*^Tg^ (Fig. 1a). In addition, increased protein levels of RRM2 and CHK1 in MEFs carrying the *Rrm2* or *Chk1* transgene were confirmed by Western blotting (Fig. 1b).

**Figure 1.**
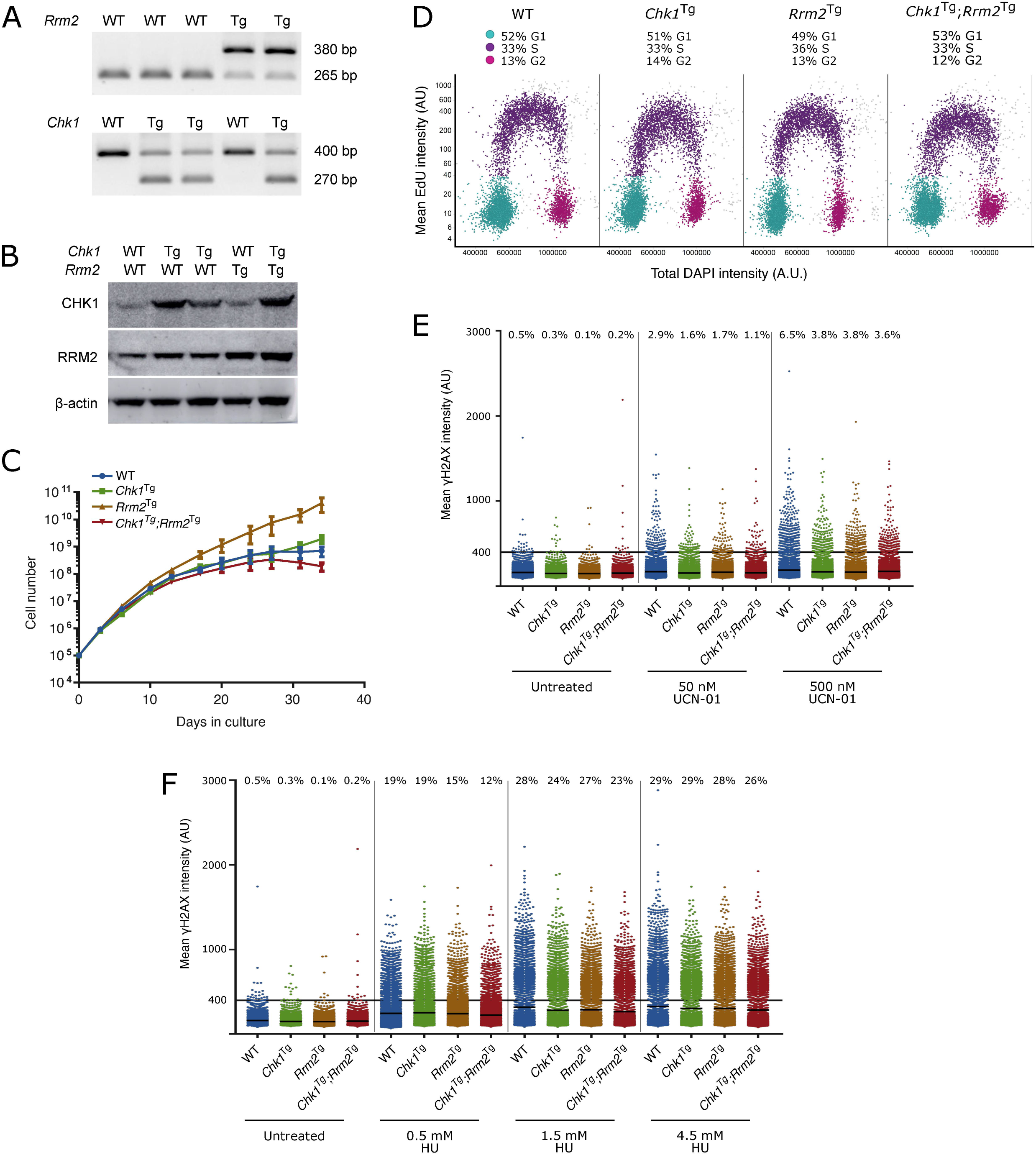
*Chk1*^Tg^, *Rrm2*^Tg^ and *Chk1*^Tg^;*Rrm2*^Tg^ MEFs are protected against replication stress. (**A**) DNA genotyping results for *Chk1* and *Rrm2* alleles in MEFs; (**B**) Western blot showing CHK1 and RRM2 protein levels in MEFs; (**C**) Proliferation curves for *Chk1* and *Rrm2* transgenic MEFs. Cells were counted and replated every 3-4 days, in three technical replicates; (**D**) Cell cycle distribution of MEFs determined by EdU incorporation and DAPI profiles. At least 7000 cells were quantified per condition using high-content microscopy; (**E**,**F**) Quantification of γH2AX intensity in MEFs treated with UCN-01 (E) or HU (F) at indicated concentrations for four hours. At least 7000 cells were quantified per condition using high-content microscopy. Percentages indicate cells with γH2AX intensity above a threshold of 400 AU, and means are indicated by horizontal black lines for each condition. The control cells are the same for (E) and (F), as the results were obtained from the same experiment.

We then assessed whether elevated levels of CHK1 and RRM2 would influence cell proliferation, and found that *Chk1*^Tg^ and *Chk1*^Tg^;*Rrm2*^Tg^ MEFs proliferated similarly to WT MEFs (Fig. 1c). *Rrm2*^Tg^ MEFs showed a mild increased proliferation compared to the other MEFs, which was consistent with an EdU staining revealing more S phase cells in the *Rrm2*^Tg^ population compared to WT, *Chk1*^Tg^ and *Chk1*^Tg^;*Rrm2*^Tg^ cells (Fig. 1d).

Next, we assessed whether MEFs carrying extra copies of *Rrm2* and *Chk1* were protected against RS by assessing γH2AX levels in these cells. Using high-content microscopy, we found that *Chk1*^Tg^, *Rrm2*^Tg^ and *Chk1*^Tg^;*Rrm2*^Tg^ MEFs have lower basal levels of γH2AX compared to WT MEFs (Fig. 1e,f). In addition to assessing γH2AX levels in unstressed conditions, we induced RS by treating the cells with CHK1-inhibitor UCN-01 at different doses and found that the cells with higher CHK1 and/or RRM2 levels showed a decrease in γH2AX intensity compared to WT MEFs (Fig. 1e). In addition, the MEFs were treated with HU (Fig. 1f). In accordance with previously published data on *Chk1* and *Rrm2* transgenic MEFs, we observed lower γH2AX intensity in these MEFs compared to WT. More importantly, the lowest levels of γH2AX intensity were observed in *Chk1*^Tg^;*Rrm2*^Tg^ MEFs. These data confirm that cells from mice carrying extra copies of the *Rrm2* or *Chk1* genes are protected from RS.

### Supraphysiological levels of CHK1 and RRM2 do not influence lifespan in mice

Next, we aimed to investigate whether the protection against RS conferred by extra copies of *Chk1* and *Rrm2* would be reflected in the survival of mice, as they did in the context of reduced levels of ATR [24,25]. To this end, we used mice containing the *Chk1* and/or *Rrm2* transgenes, and assessed tumor-free survival of these mice. Mice were euthanized when they had noticeable tumors or were visibly ill, as observed by rapid weight loss, hunched posture, rough hair coat, labored breathing, lethargy, impaired mobility or abdominal swelling. Only female mice were included in the cohorts, since they could be group-housed more easily than males.

Our results show that survival was not increased due to CHK1 or RRM2 levels: the survival of WT, *Chk1*^Tg^, *Rrm2*^Tg^ and *Chk1*^Tg^;*Rrm2*^Tg^ was not significantly different (log-rank p=0.3944) (Fig. 2a). We collected tissues of two year-old mice (not included in survival curve). H&E staining of spleens, livers and kidneys of mice did not show obvious differences between the genotypes (Fig. 2b). Furthermore, we observed no noticeable differences in appearance among the mice with different genotypes, as reflected by their weight at 1 year of age (Fig. 2c). In addition, all mice displayed comparable signs of aging such as weight loss, grey hair and baldness (data not shown). In summary, unlike in an ATR-deficient background, extra copies of *Chk1* and *Rrm2* did not increase the overall survival of mice.

**Figure 2.**
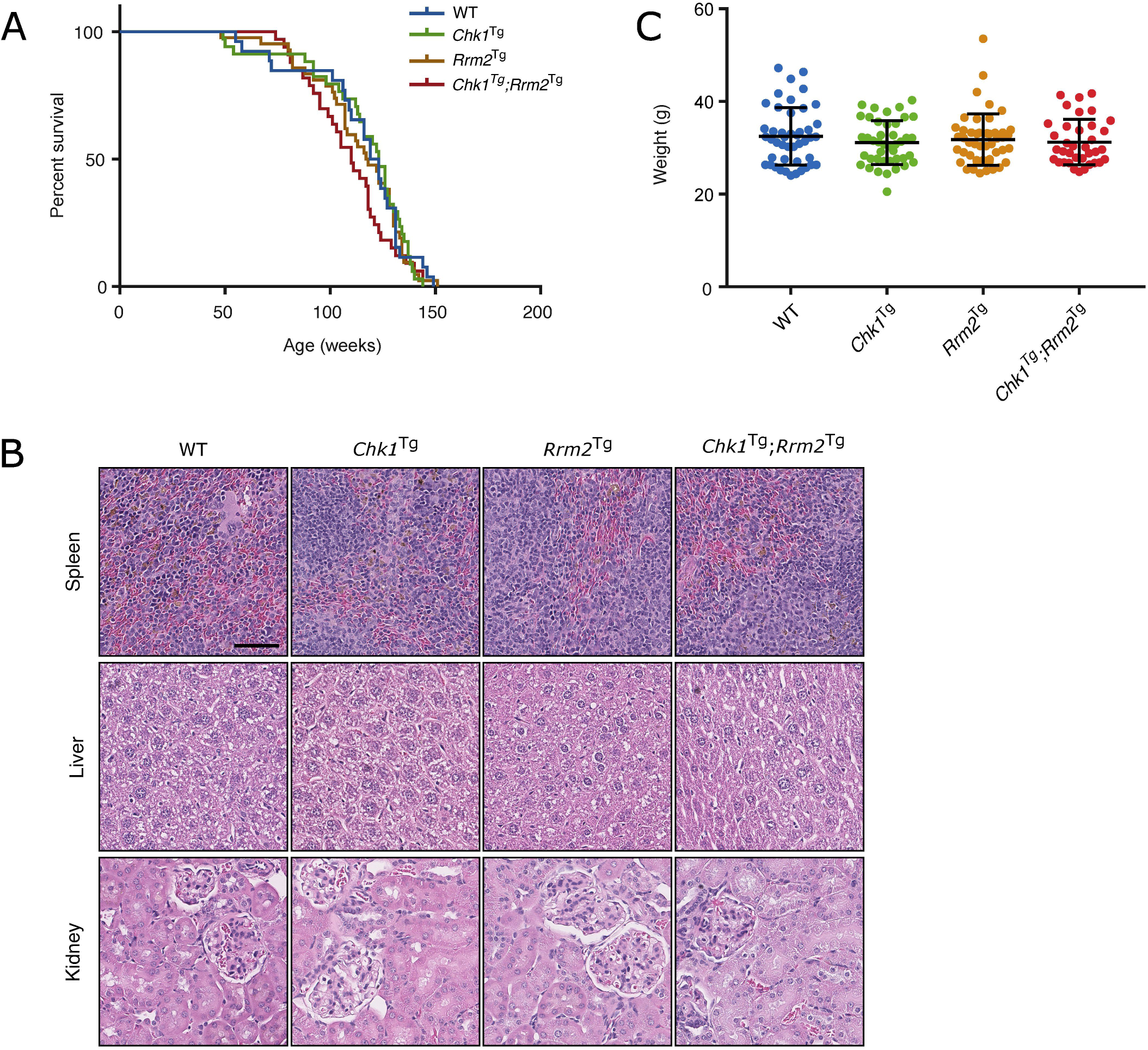
*Chk1* and *Rrm2* transgenes do not increase the lifespan of mice. (**A**) Kaplan-Meier survival curves for WT (n=26), *Chk1*^Tg^ (n=34), *Rrm2*^Tg^ (n=42) and *Chk1*^Tg^;*Rrm2*^Tg^ (n=33) mice. P-value = 0.3944 using log-rank test; (**B**) Total body weight of mice at one year of age; (**C**) Hematoxylin and eosin stainings of mouse spleen, liver and kidney for mice with the indicated genotypes. Scale bar indicates 60 µm.

### Extra copies of *Chk1* and *Rrm2* mildly increase the prevalence of spontaneous tumors

While none of the genetic combinations used in this study was able to extend mouse lifespan, the median survival of *Chk1*^Tg^;*Rrm2*^Tg^ mice was the lowest with 100 weeks compared to 123 weeks for WT mice (not statistically significant). In addition, and despite no significant differences on overall survival, the percentage of mice that developed tumors was higher for the mice containing extra copies of *Chk1, Rrm2* or both transgenes (74%, 70% and 71%, respectively) compared to the percentage of WT mice with tumor (64%), which could be in line with previous reports indicating that overexpression of CHK1 or RRM2 favors tumorigenesis (Fig. 3a) [27,28]. The majority of the observed tumors were lymphomas, which was confirmed by an enlarged spleen or lymph nodes containing CD3e positive cells (Fig. 3b). This is in agreement with our previous reports showing that increased CHK1 or RRM2 levels can reduce the DNA damage induced by oncogenes thereby facilitating oncogenic transformation or cell reprogramming [25,29,30].

**Figure 3.**
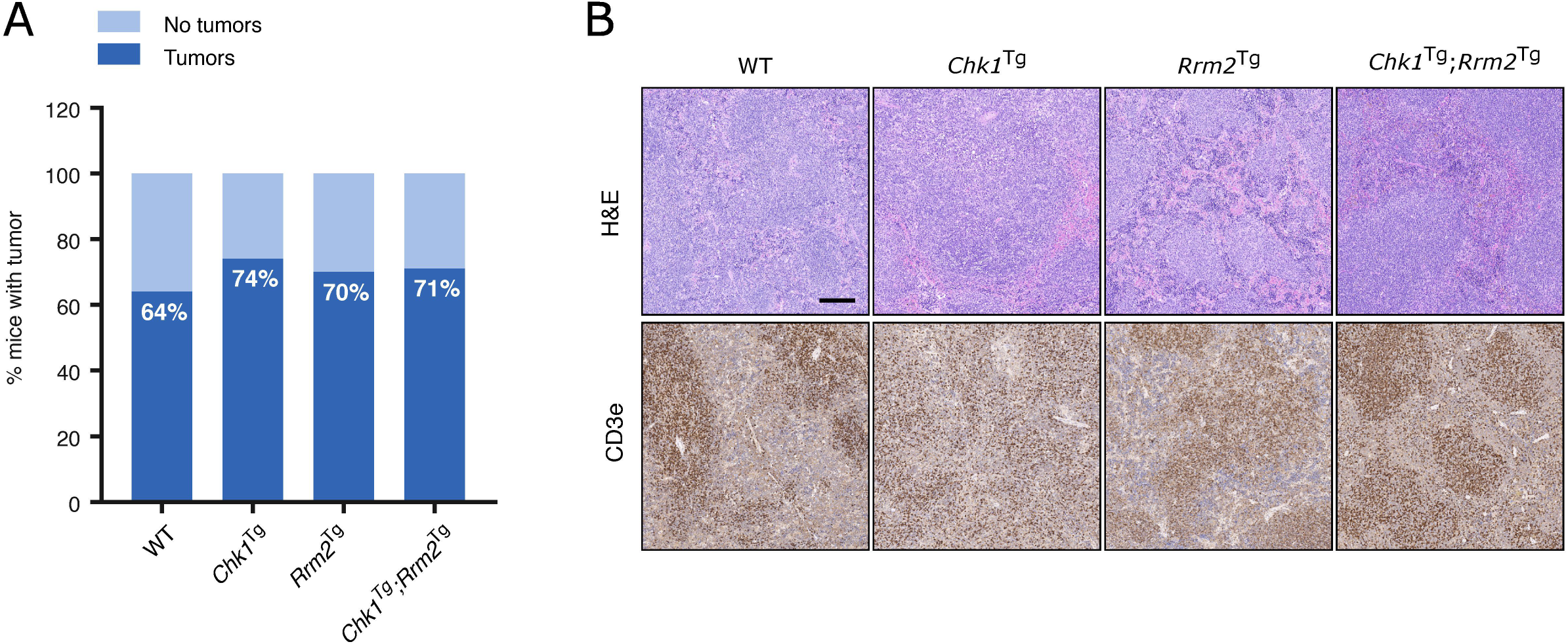
Slightly increased incidence of spontaneous tumors in *Chk1*^Tg^, *Rrm2*^Tg^ and *Chk1*^Tg^;*Rrm2*^Tg^ mice compared to WT littermates. (**A**) Tumor incidence in WT (n=22), *Chk1*^Tg^ (n=19), *Rrm2*^Tg^ (n=27) and *Chk1*^Tg^;*Rrm2*^Tg^ (n=24) mice subjected to necropsy. (**B**) H&E and CD3e IHC staining of mouse spleens with tumor found with necropsy. Tumors are CD3e positive. Scale bar indicates 200 µm.

## Discussion

In the presented study, we aimed to determine whether protection against RS could extend mammalian lifespan by utilizing mouse models with supraphysiological levels of CHK1 and RRM2. These BAC-transgenic mouse models contain extra copies of the genes of interest, which keep the expression of the genes in a physiological range and maintain their endogenous regulation and has been a successful strategy in the past [15,26]. Previous studies have shown that defects in RS-related proteins lead to increased RS levels and accelerated aging [12,31,32]. Lopez-Otin and colleagues described in 2013 the hallmarks of aging [3], one of them being DNA damage. However, a hallmark of aging should not only cause accelerated aging when aggravated, but also increase lifespan when attenuated [3].

We showed that CHK1- and RRM2-overexpressing MEFs have lower levels of DNA damage in basal conditions and have greater protection against HU- and UCN-01-induced RS, and we hypothesized that this protection against RS could lead to an increase in lifespan *in vivo*. However, we did not observe a difference in lifespan between *Chk1*^Tg^, *Rrm2*^Tg^ and *Chk1*^Tg^;*Rrm2*^Tg^ and WT mice, and all mice developed similar age-related symptoms. Based on these results we conclude that supraphysiological levels of CHK1 and RRM2 do not influence normal aging, and more generally, we propose that physiological RS might not be an important driver of aging. Although *Chk1*^Tg^ and *Rrm2*^Tg^ mice were previously shown to extend the lifespan and alleviate the progeroid symptoms of ATR-deficient mice [24,27], this is not a direct indication that CHK1 and RRM2 levels influence normal aging. CHK1 overexpression could compensate for ATR deficiency, as CHK1 is a downstream target of ATR. For RRM2 this connection to the ATR pathway is not as direct, although studies in yeast have shown that ATR ortholog Mec1 can activate the ribonucleotide reductase (RNR) complex [33,34]. Therefore, extra levels of RRM2 could also partially compensate for ATR deficiency.

Interestingly, a slightly higher fraction of the *Chk1* and *Rrm2* transgenic mice developed tumors, primarily lymphomas, compared to WT mice. The exact role of the RS response in regard to tumorigenesis is complex [35,36]. On the one hand, CHK1 is known as a tumor suppressor, and its expression can induce senescence and apoptosis, thereby limiting tumorigenesis [37,38]. On the other hand, experiments with *Chk1*^Tg^ MEFs have shown that overexpression of CHK1 can promote oncogenic transformation, and elevated CHK1 levels are present in lymphomas, suggesting that CHK1 could also have a role in promoting tumorigenesis [27,39]. In relation to RRM2, nucleoside supplementation has been shown to both limit and promote transformation [40,41]. In addition, increased nucleotide levels can lead to errors during DNA replication and tumorigenesis [28,42]. Our findings favor the concept that RS is not a requirement for transformation, but also that protection against RS does not prevent tumorigenesis.

Furthermore, the increased tumor incidence in *Chk1* and *Rrm2* transgenic mice could affect their survival, as is the case for TERT-overexpressing mice. Several *Tert* trangenic mouse models have an increased incidence of spontaneous tumors [43–46]. Thus, extension of lifespan by TERT-overexpression is only possible in the context of a cancer-resistant background [16]. Since we observed a slightly higher tumor incidence in *Chk1* and *Rrm2* transgenic mice, these mice could possibly survive longer in a cancer-resistant background. Also, the process of aging is complex and influenced by different factors. Conversely, overexpression of DNA damage proteins may improve several aspects of aging, but not the aging process as a whole, and therefore we cannot rule out the involvement of RS on aging. Nevertheless, our data indicate that RS might not be a relevant contributing factor to normal aging in mice.

## Methods

### Mouse husbandry

*Chk1*^Tg^ and *Rrm2*^Tg^ mouse models have been described previously [24,25]. Both strains have a mixed C57BL/6-129/Sv background and carry a bacterial artificial chromosome (BAC) transgene. In case of *Chk1*^Tg^ mice, there is one extra copy of the gene; *Rrm2*^Tg^ mice carry multiple copies of *Rrm2*^Tg^ in tandem. The mice in this study were housed at the University of Copenhagen, Department of Experiment Medicine and mouse work was monitored by the Institutional Animal Care and Use Committee and performed in compliance with Danish and European regulations.

### Primers for PCR Genotyping

Mice were genotyped using the following primers: gRrm2_Fw (TGTCCTGGAGAGCCAGTCTT), gRrm2_Rev (AAGGAGGGAGGGAGGCTATT) and gRrm2_Transgen (ACTGGCCGTCGTTTTACAAC) for the *Rrm2* locus and gChk1_Fw (TGTCTTCCCTTCCCTGCTTA), gChk1_Rev (TCCCAAGGGTCAGAGATCAT) and gChk1_Transgen (GTAAGCCAGTATACACTCCGCTA) for the *Chk1* locus. Expected band sizes are 265 bp for the *Rrm2* ^+^, 380 bp for the *Rrm2*^Tg^, 400 bp for the *Chk1*^+^ and 270 bp for the *Chk1*^Tg^ allele.

### Cell culture

Mouse embryonic fibroblasts (MEFs) were generated from 13.5 dpc mouse embryos according to standard procedures. MEFs were cultured in Dulbecco’s Modified Eagle Medium (DMEM; Gibco) supplemented with 15% heat-inactivated fetal bovine serum (FBS; ThermoFisher Scientific) and 1% penicillin-streptomycin (ThermoFisher Scientific).

### Proliferation assay

Early-passage MEFs were plated at a density of 100.000 cells per well of a 6-well plate. Every three to four days, cells were counted and re-seeded at a density of 100.000 wells per well. The experiment was carried out in three technical replicates.

### Cell cycle analysis

10.000 cells were plated per well of a 96 well optical plate. After 24 hours, cells were incubated with 10 µM EdU (Life Technologies, A10044) for 30 minutes, before fixation in 4% PFA. Cells were permeabilized with 0.5% Triton X-100 in PBS for 15 min and washed three times with PBS. Click-iT reaction cocktail containing CuSO_4_, L-ascorbic acid and Alexa Fluor 647 fluorescent dye azide in PBS was added to each well and incubated according to the manufacturer’s protocol (Invitrogen C10269). Nuclei were counterstained with DAPI, and cell cycle profiles were made by image acquisition on an automated Olympus IX83 ScanR microscope and Olympus ScanR analysis software.

### DNA damage experiments and drug treatments

MEFs at passage four were seeded in a 96 well optical plate at a density of 10.000 cells/well. After 24 hours, MEFs were incubated with hydroxyurea (Sigma-Aldrich H8627) or UCN-01 (Sigma Aldrich U6508) at indicated concentrations for four hours, followed by fixation with 4% paraformaldehyde (PFA). For immunofluorescence experiments, cells were permeabilized with Triton X-100 (Sigma-Aldrich T8787), blocked in IF blocking buffer (3% BSA, 0.1% Tween in PBS), and incubated with γH2AX antibody (Merck Millipore 05-636) at 4°C overnight. Samples were incubated with goat anti-mouse IgG secondary antibody, Alexa Fluor 488 (Sigma-Aldrich A-11001) for 1 hour at room temperature, and nuclei were stained with DAPI. High-content microscopy was performed using an automated Olympus IX83 ScanR microscope.

### Western blotting

Cells were lysed in RIPA buffer (Sigma-Aldrich R0278) supplemented with protease (Roche 4693132001) and phosphatase inhibitors (Sigmal-Aldrich P0044). Western blot analysis was performed according to standard procedures. Primary antibodies used were anti-CHK1 (Novocastra), anti-RRM2 (Santa Cruz Biotechnology, SC-10844) and anti-β-actin (Sigma-Aldrich, A5441).

### Histopathology

Mouse tissues were fixed in formalin and processed for immunohistochemistry (IHC) by HistoWiz Inc. (histowiz.com) or the Histopathology unit at the Spanish National Cancer Research Center (CNIO).

## Author contributions

A.J.L.-C. conceived and supervised the study. E.A. designed and performed most of the experiments. All the authors provided relevant intellectual input during this study. M.S. and A.A helped with the maintenance of the mouse colonies and generation of MEFs. The transgenic mouse models were generated in O.F. lab. A.J.L.-C. and E.A. wrote the manuscript. All of the authors read, edited and commented on the manuscript. We thank the caretakers at Department of Experimental Medicine at the University of Copenhagen, and the Histopathology core unit at the Spanish National Cancer Research Center (CNIO, Madrid) for technical support

## Funding

Research in A.J.L.-C. lab was funded by Danish Council for Independent Research (Sapere Aude, DFF-Starting Grant 2014); Danish National Research Foundation (DNRF115); Danish Cancer Society (KBVU-2017_R167-A11063); European Research Council (ERC-2015-STG-679068); and Nordea-fonden (02-2017-1749). Research in O.F. lab was funded by grants from the Spanish Ministry of Science; Innovation and Universities (RTI2018-102204-B-I00, co-financed with European FEDER funds); and the European Research Council (ERC-617840).

## Conflict of interest statement

None declared.

